# Tau expression and phosphorylation in enteroendocrine cells

**DOI:** 10.1101/2023.02.19.528206

**Authors:** Guillaume Chapelet, Nora Béguin, Blandine Castellano, Isabelle Grit, Thibauld Oullier, Michel Neunlist, Hervé Blottière, Malvyne Rolli-Derkinderen, Gwenola Le Dréan, Pascal Derkinderen

## Abstract

**Background and objective:** There is mounting evidence to suggest that the gut-brain axis is involved in the development of Parkinson’s disease (PD). In this regard, the enteroendocrine cells (EEC), which faces the gut lumen and are connected with both enteric neurons and glial cells have received growing attention. The recent observation showing that these cells express alpha-synuclein, a presynaptic neuronal protein genetically and neuropathologically linked to PD came to reinforce the assumption that EEC might be a key component of the neural circuit between the gut lumen and the brain for the bottom-up propagation of PD pathology. Besides alpha-synuclein, tau is another key protein involved in neurodegeneration and converging evidences indicate that there is an interplay between these two proteins at both molecular and pathological levels. There are no existing studies on tau in EEC and therefore we set out to examine the isoform profile and phosphorylation state of tau in these cells.

**Methods:** Surgical specimens of human colon from control subjects were analyzed by immunohistochemistry using a panel of anti-tau antibodies together with chromogranin A and Glucagon-like peptide-1 (two EEC markers) antibodies. To investigate tau phosphorylation and expression further, two EEC lines, namely GLUTag and NCI-H716 were analyzed by western blot after dephosphorylation with pan-tau and tau isoform specific antibodies. Eventually, GLUTag were treated with propionate and butyrate, two short chain fatty acids known to sense EEC, and analyzed at different time points by western blot with an antibody specific for tau phosphorylated at Thr205.

**Results:** We found that tau is expressed and phosphorylated in EEC in adult human colon and that both EEC lines mainly express two tau isoforms that are phosphorylated under basal condition. Both propionate and butyrate regulated tau phosphorylation state by decreasing its phosphorylation at Thr205.

**Conclusion and inference:** Our study is the first to characterize tau in human EEC and in EEC lines. As a whole, our findings provide a basis to unravel the functions of tau in EEC and to further investigate the possibility of pathological changes in tauopathies and synucleinopathies.

## Introduction

The enteroendocrine cells (EEC), which are scatterly distributed along the entire gastrointestinal (GI) mucosa representing around 1% of the total gut epithelium cell population, are key components of the gut-brain axis. They are classically regarded as specialized hormone-secreting cells with an apical surface that is exposed to gut lumen and a basal portion that contains secretory granules (Gribble and Reimann, 2019). Such an orientation allows EEC to respond to intraluminal signals such as nutrients or microbiota-derived metabolites. This aspect has been particularly well documented for short-chain fatty acids (SCFA), such as butyrate and propionate, which induce the release of the peptide hormone YY (PYY) (Larraufie et al., 2018). Morphologically, EEC were shown to exhibit neuron-like features including the expression of pre- and post-synaptic proteins together with the presence of neurite-like processes, called neuropods (Bohórquez et al., 2015). These neuropods are in close contact with the two components of the enteric nervous system (ENS), namely enteric glial cells (Bohórquez et al., 2014) and enteric neurons (Chandra et al., 2017) as well as vagal neurons (Kaelberer et al., 2018). The recent findings, which showed that EEC contain alpha-synuclein (Chandra et al., 2017), a presynaptic neuronal protein genetically and neuropathologically linked to Parkinson’s disease (PD) led to the assumption that they might be involved in the development of PD. By facing the gut lumen and being directly connected with alpha-synuclein positive enteric neurons (Chandra et al., 2017), the EEC might be a key component of the neural circuit between the gut lumen and the brain for the bottom-up propagation of PD pathology as initially hypothesized by Braak (Braak et al., 2006) and more recently by Borghammer (Borghammer and Van Den Berge, 2019).

Alpha-synuclein accumulates in a group of neurodegenerative diseases collectively known as synucleinopathies with PD being the most common, while the accumulation of tau is a defining feature of tauopathies classically found in the brains of patients with Alzheimer’s disease (AD) or progressive supranuclear palsy (PSP) (Dugger and Dickson, 2017). Nevertheless, in numerous cases alpha-synuclein positive inclusions are also described in tauopathies and *vice versa*, suggesting a co-existence or crosstalk of these proteinopathies (Arai et al., 2001; Coughlin et al., 2019). Tau, like alpha-synuclein is expressed by enteric neurons (Lionnet et al., 2018), thereby suggesting that enteric tau might be involved in neurodegenerative disorders and/or enteric neuropathies (Derkinderen et al., 2021). In contrast to alpha-synuclein, no data are available about the distribution and phosphorylation pattern of tau isoforms in EEC. Here, we first examined the expression levels of tau isoforms and their phosphorylation profile in full thickness segments of human colon and in EEC lines. We then studied the regulation of tau phosphorylation by SCFA in EEC. Our results show the presence of phosphorylated tau in human colonic EEC and the expression of two main human tau isoforms in EEC lines. EEC tau is phosphorylated and its phosphorylation state can be modified by SCFA. These data provide the first detailed characterization of EEC tau in human adult colonic tissues and in cell lines. Further investigation of tau modifications in EEC in pathological conditions may provide valuable information about the possible role of EEC tau in neurodegenerative diseases.

## Material and methods

### Human and mouse tissues

Specimens of human colon were obtained from 8 neurologically unimpaired subjects who underwent colon resection for colorectal cancer (5 men, 71±7.6 years). For all tissues specimens, sampling was performed in macroscopically normal segments of uninvolved resection margins. The sampling of human colon was approved by the Fédération des biothèques of the University Hospital of Nantes, according to the guidelines of the French Ethics Committee for Research on Humans and registered under the no. DC-2008-402. Written informed consent was obtained from each subject. Hippocampi from a 2-month-old mouse were removed and stored at -80 °C until further analysis by western blot.

### EEC lines and reagents

NCI-H716 cells (ATCC, LGC Standards, Molsheim, France, CCL-251) and GLUTag cells (a gift from Colette Roche, INSERM Lyon) were maintained respectively in RPMI-1640 (Gibco, Life Technologies, Villebon-sur-Yvette, France) and in DMEM (Merck-Sigma, Molsheim France), all supplemented with 10% fetal bovine serum, 2 mM L-Glutamine, 50 IU/mL penicillin and 50 μg/mL streptomycin (all from Merck-Sigma) in a humidified incubator at 37 °C with 5% of CO_2_. Propionate and butyrate were from Merck-Sigma.

### Dephosphorylation of tissues and cell lysates

For dephosphorylation experiments, cells and hippocampi were homogenized in a buffer containing 100 mM NaCl and 50 mM Tris-Cl at pH 7.4 with 1% (v/v) IGEPAL® CA-630 (ThermoFisher, Saint-Herblain, France) and a protease inhibitors cocktail without EDTA (Roche, Neuilly sur Seine, France) using either a “Precellys 24” (Bertin technologies, St Quentin-en-Yvelines, France) tissue homogenizer and followed by sonication with “vibracell 75 186” device (Sonics, Newton CT, USA). Homogenates were centrifuged at 16,100 g for 20 min at 4 °C with an Eppendorf 5415R centrifuge (Eppendorf, Hamburg, Germany), sonicated for 10 s and protein amounts normalized following a bicinchoninic acid protein assay (ThermoFisher). Samples were diluted to 1.0 mg/mL protein using homogenization buffer and incubated with 20 U/μL lambda phosphatase in MnCl_2_ and enzyme buffer as supplied with the lambda protein phosphatase kit (New England Biolabs, Evry-Courcouronnes, France) for 3 h at 30 °C. The reaction was stopped by the addition of sample buffer (Life Technologies) and heating to 95 °C for 5 min. Control samples were treated identically without the addition of lambda phosphatase.

### SDS-PAGE and western blot

For dephosphorylation experiments, cells or tissues were processed as described above. For experiments that did not require dephosphorylation, cells or tissues were lysed in RIPA lysis buffer (Merck Millipore, Fontenay sous Bois, France) containing 2 mM orthovanadate (Merck-Sigma, Molsheim France), phosphatase inhibitor cocktail II (Merck-Sigma) and a protease inhibitors cocktail (Sigma). Western blots were performed as we previously described using NuPAGE™ 10% Bis-Tris Protein Gels (Life Technologies). The primary anti-tau antibodies used are listed in Table 1. β-actin antibodies (Abcam, France, 1:1000 dilution) were used for as loading control. Tau ladder (six human tau recombinant isoforms, Sigma-Merck) was used to identify tau isoforms in NCI-H716 cells. For quantification, the relevant immunoreactive bands were quantified with laser-scanning densitometry and analyzed with Image Lab software (Biorad, Marnes-la-Coquette, France) and image J software (NIH; version 1.51). To allow comparison between different films, the density of the bands was expressed as a percentage of the average of controls. pThr205 tau immunoreactive bands were measured, normalized to the optical densities of total tau, and expressed as percentage of controls.

**Table 1.**
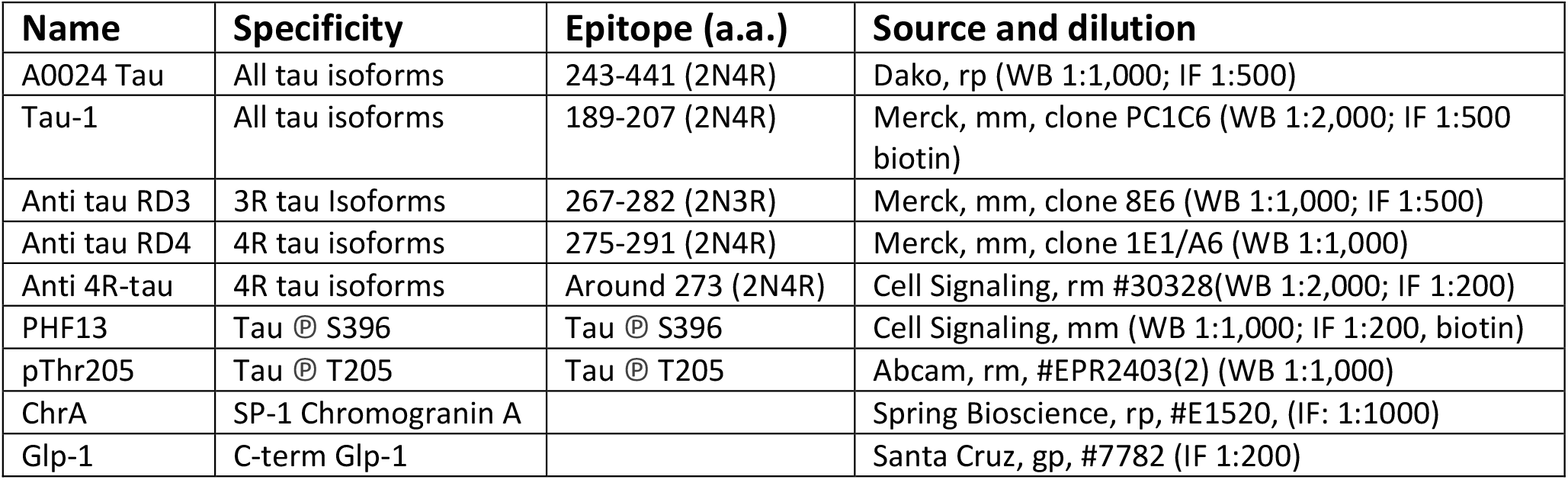
The name, specificity, epitope, source and dilution of the antibodies used in this study are shown. Abbreviations are amino-acids (a.a.); immunofluorescence (IF); goat polyclonal (gp); mouse monoclonal (mm); rabbit monoclonal (rm); rabbit polyclonal (rp); western blot (WB).

### Immunofluorescence

Human colonic tissues were embedded in paraffin using an embedding station (LEICA EG1150C) and sections (3 µm) were cut using a microtome (LEICA RM2255). The sections were deparaffinized by bathing twice in xylene (for 5 min each) and taken through graded concentrations of ethanol (100%, 95%, 70%, 50%, respectively for 3 min each). After a rinse in distilled water, slides were washed in PBS and antigen retrieval was performed using a Sodium Citrate solution (2.94 g Sodium Citrate Tribase ; 1L distilled water ; 500 µL Tween 20 ; pH 6) at 95°C for 20 min. Slides were incubated in NH_4_Cl (100 mM) for 15 min before incubation in PBS-0.5% triton X-100 for 1 hour and blocking for 2 hours in 10 % (v/v) horse serum in PBS-0.5% triton X-100. Primary antibodies (Table 1) were incubated overnight at 4°C, and following washing, secondary antibodies were added for 2 hours at room temperature. Secondary antibodies were anti-mouse Biotin (A24522, ThermoFisher scientific), Alexa Fluor 647-conjugated goat anti-rabbit (711-606-152, Jackson Immunoresearch, Ely, UK), Alexa Fluor 568-conjugated Streptavidin (S11226, Invitrogen, ThermoFisher scientific), Alexa Fluor 568-conjugated goat anti-mouse (Molecular Probes, Thermo Scientific). DAPI (1:10,000) was added to counterstain nuclei. Tissues sections were mounted in Prolong Gold anti-fading medium (Molecular Probes, Thermo Scientific). Images were acquired with a Zeiss, Axio Imager M2m fluorescence microscope coupled to a digital camera (Axiocam 503 mono).

### Statistics

All quantitative data shown are mean ± standard error of the mean (SEM). Mann-Whitney test or Kruskal-Wallis then Dunn’s multiple comparisons tests were performed for two or multiple groups comparison, respectively. Differences were deemed statistically significant if p < 0.05. GraphPad Prism software version 9.1.1 (GraphPad Software Inc., La Jolla, California, United States) was used for statistical analyses and for designing figures.

## Results

### Human colonic EEC express tau

To determine if tau is expressed in EEC, we examined chromogranin A and glucagon-like peptide 1 (GLP-1)-containing cells of the human colon. Chromogranin A is an acidic glycoprotein located in secretory vesicles of endocrine cells, which is classically used as a pan-EEC marker (Wilson and Lloyd, 1984), whereas GLP-1 is a marker of L-type EEC (Eissele et al., 1992), the most abundant EEC population in the human colon (Latorre et al., 2016). Using the pan-tau antibody A0024, we identified tau in GLP-1-positive cells in the epithelial lining (Fig 1a). When human colonic tissue was immunolabelled with isoform-specific tau antibodies, we showed that 3R-tau was expressed in chromogranin A immunoreactive cells (Fig 1b); no specific staining was observed when a 4R-tau was used (data not shown). Tau phosphorylation at multiple serine and threonine sites is the predominant mechanism by which its biological activity is regulated (Guo et al., 2017). We therefore examined the phosphorylation state of tau with a phospho-specific antibody that detect tau phosphorylated at Ser396 and showed that phospho-tau was observed within chromogranin A-positive cells in colon epithelium (Fig 1c). When taken together, these results show that tau is expressed and phosphorylated in EEC in human colon, with 3R-tau likely being most abundant isoforms.

**Figure 1.**
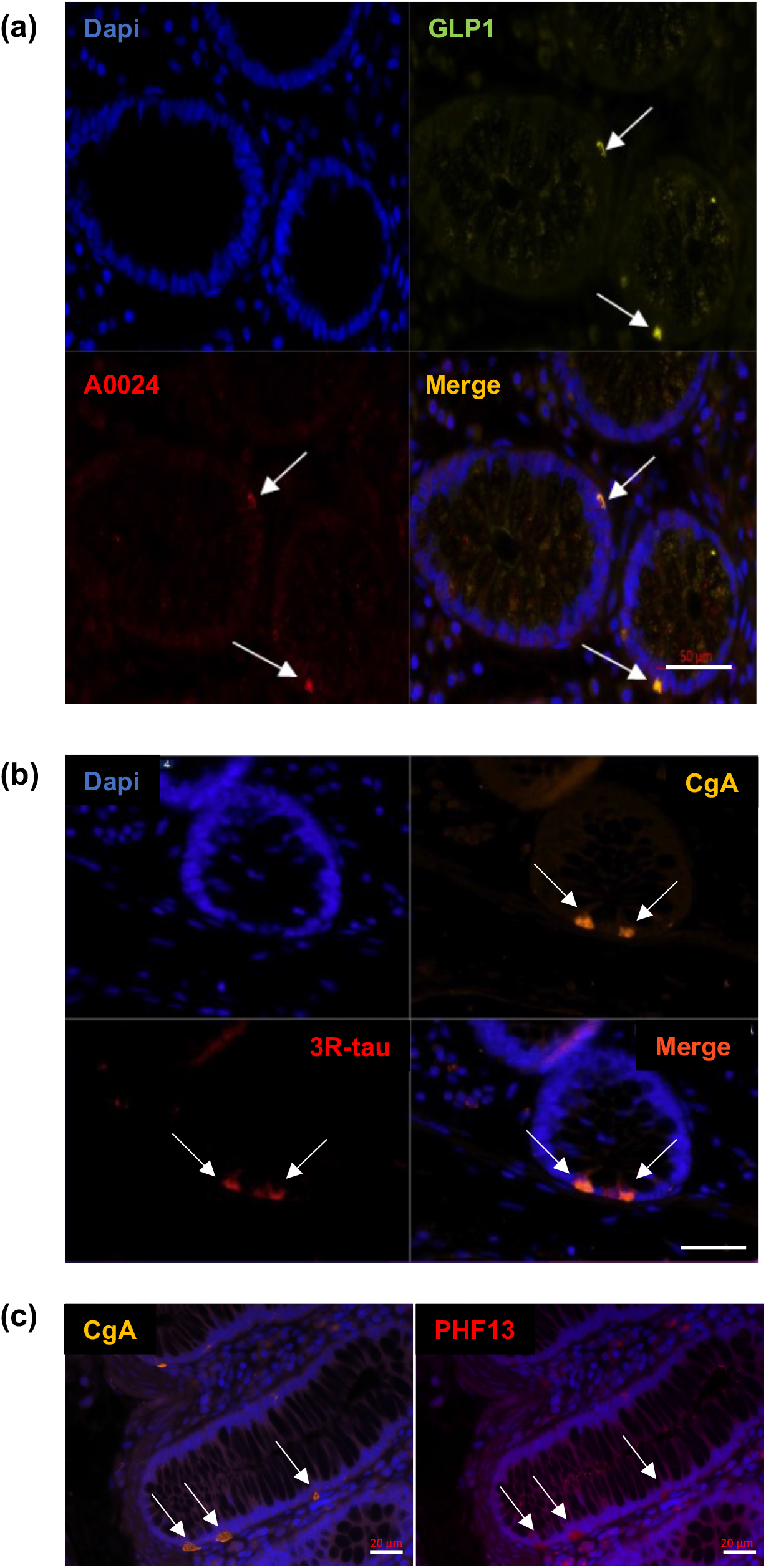
Distribution of tau in the epithelial lining in adult human colon. **(a)** Total tau antibody A0024 (table 1) was used to detect tau in epithelium lining of human colonic samples (arrows). An antibody specific to Glucagon-like peptide 1 (GLP-1) was used to specifically label L-type EEC (arrows). DAPI was used to counterstain nuclei. Scale bar is 50 μm **(b)** Isoform-specific antibody (tau RD3, table 1) was used to detect 3R-tau and an antibody specific to chromogranin A (CgA) was used to specifically label EEC (arrows). DAPI was used to counterstain nuclei. Scale bar is 50 μm **(c)** Human colonic samples were co-labelled with an antibody specific for tau phosphorylated at Ser396 (PHF13, table 1) and an antibody specific to chromogranin A (CgA). Arrows show cells immunoreactive for both PS396 and CgA. DAPI was used to counterstain nuclei. Scale bar is 20 μm. Representative photomicrographs are shown.

### Tau is expressed and phosphorylated in EEC lines

There are multiple cell lines used as models for EEC research, including GLUTag, a mouse endocrine tumor-adherent cell line and NCI-H716, a human-derived suspension cell line. These two cell lines have been widely used as models for GLP-1–producing L-cells and proved to be useful for *in vitro* screening bioassays (Goldspink et al., 2018). As a first approach to identify tau isoforms in EEC lines, we compared the banding pattern on western blots of total tau as evaluated with the A0024 pan-Tau antibody between GLUTag cells and hippocampus of 2-month-old mouse. In keeping with previous findings, total tau antibody detected a tau quadruplet in 2-month old mouse brain with three major bands between 50 and 60 kDa and a fainter one around 48 kDa, which likely correspond to 0N4R-1N4R-2N4R and 0N3R isoforms, respectively (McMillan et al., 2008; Liu and Götz, 2013) (Figure 2a). In GLUTag cells, the observed banding pattern was markedly different with a doublet of 48 and 50 kDa bands, the latter showing the most intense labelling (Figure 2a). In adult mouse brain, primary CNS/ENS neurons and neuronal cell lines, tau isoforms are phosphorylated on multiple tau and serine residues resulting in reduced electrophoretic mobility on SDS-PAGE compared to non-phosphorylated tau (Goedert et al., 1992; Davis et al., 1995; Lionnet et al., 2018). In order to determine the phosphorylation state of tau in GLUTag cells, cell lysates treated or not with lambda phosphatase (Davis et al., 1995) were analyzed by western blot using 3 different pan-tau antibodies. Treatment with lambda phosphatase caused tau dephosphorylation, as evidenced by a significant downward shift in mobility of the tau doublet detected with either the pan-Tau A0024, D1M9X or Tau-1 (Figure 2b). Of note, this downward shift was associated with increased Tau immunoreactivity when the Tau-1 antibody was used (Figure 2b). These findings are in line with previous observations showing that Tau-1 binds preferentially to tau when dephosphorylated at serine residues 195, 198, 199, and 202 (Liu et al., 1993; Szendrei et al., 1993). To further refine the analysis of tau isoforms in GLUTag cells, we used 2 commercially available isoform-specific tau antibodies directed against 3R and 4R-tau, which have been shown to be highly specific in a recent comprehensive study that tested the specificity of tau antibodies using immunoblotting (Ercan et al., 2017). GLUTag lysates were compared to dephosphorylated brain samples (hippocampus of 2-month old mice), which, as mentioned above, express all 4R isoforms and to a lesser extent the 0N3R isoform (McMillan et al., 2008; Liu and Götz, 2013) (Figure 2c). After dephosphorylation, the 3R and 4R antibodies detected one single band in cell lysates that comigrated with 0N3R and 0N4R in the hippocampus, respectively (McMillan et al., 2008; Liu and Götz, 2013)(Figure 2c). When taken together, these results show that 0N3R and 0N4R are the two main tau isoforms that are expressed in GLUTag cells and these two isoforms are phosphorylated under basal conditions. Similar experiments were conducted with the NCI-H716 cell line. Treatment with lambda phosphatase caused tau dephosphorylation, as evidenced by a significant downward shift in mobility of the tau doublet detected when the pan-Tau A0024 antibody was used (Figure 2d). After dephosphorylation, this tau doublet at 50 and 55 kDa comigrated with the 0N3R and 1N3R/0N4R from the tau human ladder (Figure 2d). The isoform specific 3R antibody detected one major at band at 58 kDa and a fainter one around 55 kDa in dephosphorylated samples that comigrated with 1N3R and 0N3R isoforms of recombinant human tau, respectively (Figure 2d). No major bands were observed when the 4R-tau antibody was used (data not shown). As a whole, these results show that 1N3R and 0N3R are the two main isoforms expressed in NCI-H716 cell.

**Figure 2.**
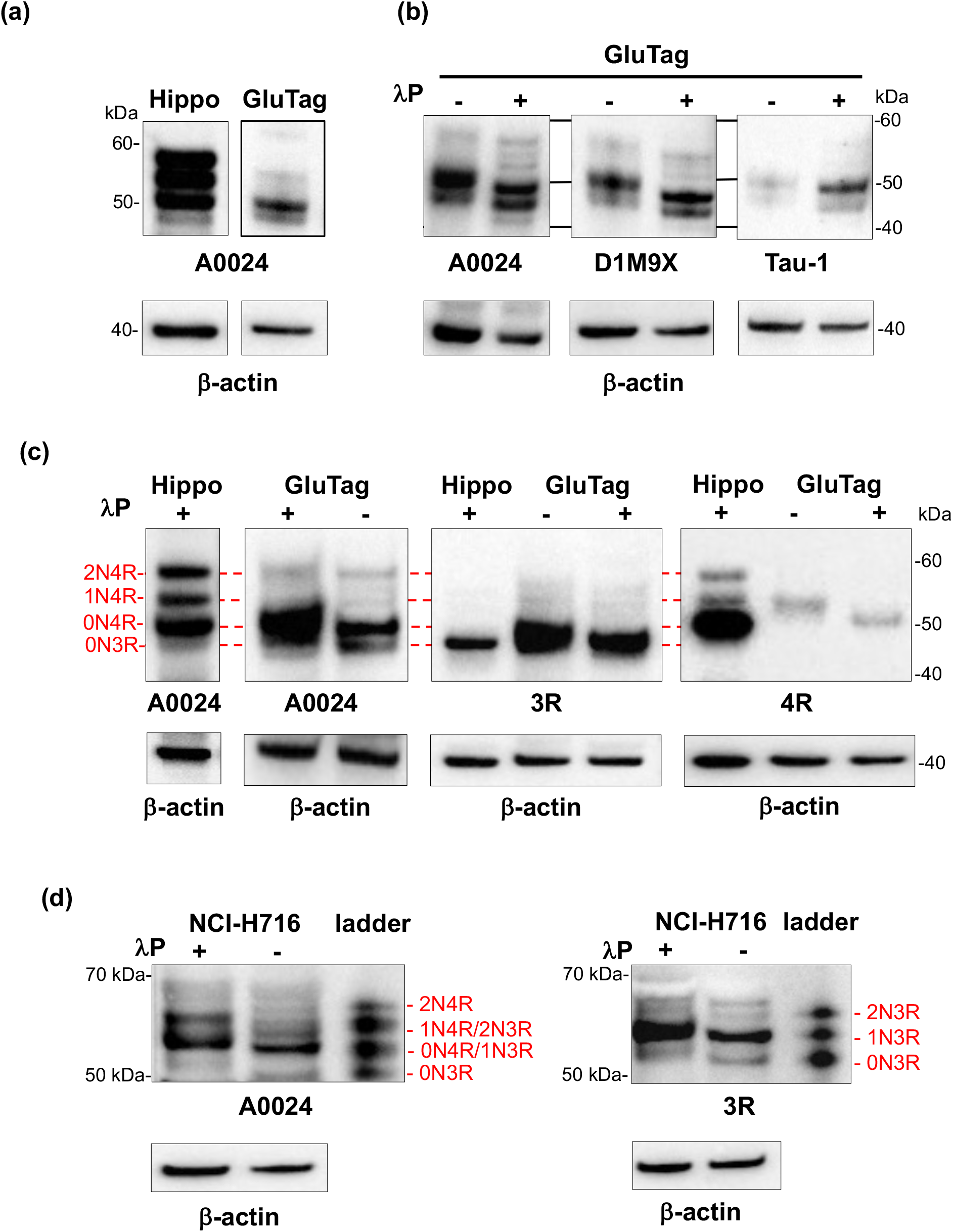
Tau isoforms and phosphorylation in EEC lines. **(a)** Hippocampus (2-month-old mice, Hippo) and GLUTag cell lysates were subjected to immunoblot analysis using the pan-Tau antibody A0024 **(b)** Lysates of GLUTag cell were treated (+) or not (−) with lambda phosphatase (λP) before immunoblotting with the pan-tau antibodies A0024, D1M9X and Tau-1 **(c)** GLUTag cells lysates were treated (+) or not (−) with lambda phosphatase (λP) before immunoblotting with the tau isoform-specific antibodies 3R and 4R and the pan-tau antibody A0024. Hippocampus lysate from 2-month old mice dephosphorylated with lambda phosphatase (λP+), which contains all 4R isoforms and 0N3R isoform (McMillan et al., 2008; Liu and Götz, 2013) was used as a ladder to determine mouse tau isoform profile in GLUTag cells; the red lines show comigration **(d)** NCI-H716 cell lysates were treated (+) or not (−) with lambda phosphatase (λP) before immunoblotting with the tau isoform-specific antibody 3R and the pan-tau antibody A0024. Tau ladder, which contains all 6 dephosphorylated human tau isoforms. In all experiments, β-actin immunoblot was used as a loading control. The results shown in (a) and (c) are representative of 3 and 2 independent experiments, respectively. The results shown in (b) are representative of 7, 2 and 3 independent experiments for A0024, DM19X and Tau-1 antibodies, respectively.

### Tau phosphorylation is regulated by SCFA in EEC lines

SCFA act as ligands for G protein coupled receptors, which are expressed in EEC, where they mediate hormone release, such as PYY and GLP-1 (reviewed in (Martin-Gallausiaux et al., 2021)). The phosphorylation of tau at multiple serine and threonine sites has been described in both developing and adult brain and is the predominant mechanism by which tau functions are regulated (Guo et al., 2017). This logically led us to study the effects of SCFA on tau phosphorylation in EEC. To this end, GLUTag cell lines were treated with 1 mM of either propionate or butyrate at different time points from 10 to 180 minutes and cell lysates were analyzed by western blot with an antibody specific for tau phosphorylated at Thr205. Treatment with propionate or butyrate for 180 min caused tau dephosphorylation at Thr205 when compared to control conditions (Figure 3 a,b), without significant changes in tau expression (Figure 3 a,b).

**Figure 3.**
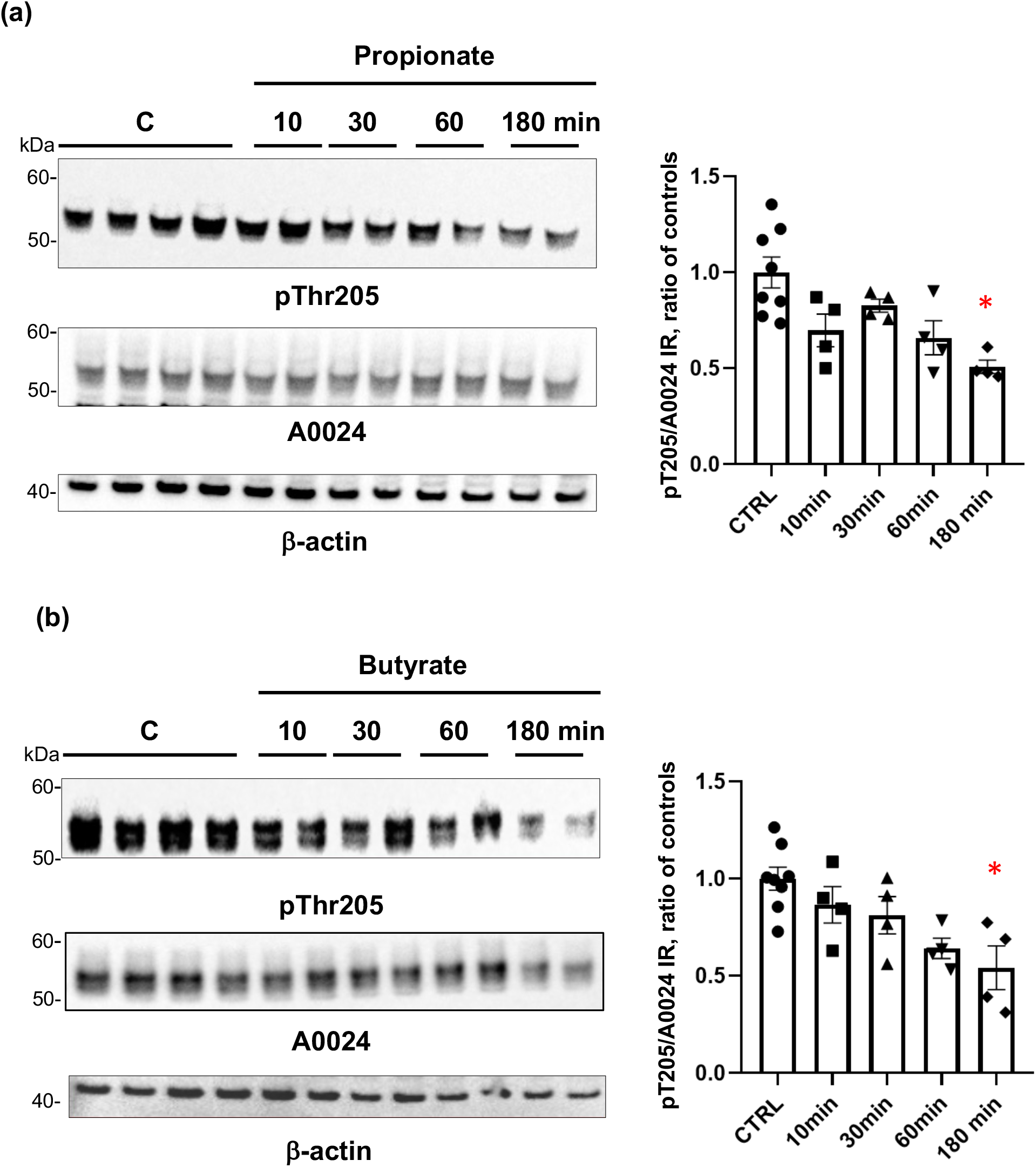
Regulation of tau phosphorylation by SCFA in GLUTag cells. **(a)** GLUTag cells were treated with 1 mM propionate for the indicated periods of time. Cell lysates were subjected to immunoblot analysis using antibodies against tau phosphorylated at Thr205 (pThr205) and total tau (A0024). Membranes were also probed with an anti-β-actin antibody to ensure equal protein loading. pThr205 immunoreactive bands were measured, normalized to the optical densities of total tau (A0024), and expressed as ratio of controls. Data correspond to mean ± SEM (n = 4-8, n indicates the number of wells; *p<0.05, 180 min-treated vs control) **(b)** GLUTag cells were treated with 1 mM butyrate for the indicated periods of time. Cell lysates were analyzed as in (a) and quantification was performed as in (a). Data correspond to mean ± SEM (n = 4-8, n indicates the number of wells; *p<0.05, 180 min-treated vs control)

## Discussion

Tau is a microtubule-associated protein for which the physiological functions are still a topic of intense investigation (Guo et al., 2017). Additionally, in pathological conditions tau is a key player in the pathogenesis of several diseases collectively referred to as tauopathies including AD and PSP (Guo et al., 2017). In the adult brain, tau is classically described as a neuronal protein specifically localized and highly enriched in axons but precise localization studies showed that tau distribution in the mature CNS is more widespread than initially thought, with expression in the somatodendritic compartment of neurons as well as in glial cells (Kanaan and Grabinski, 2021). Besides the CNS, the presence of tau has been demonstrated in several non-neuronal cells, such as monocytes (Kim et al., 1991), lymphocytes (Kvetnoy et al., 2000), testicular spermatids (Ashman et al., 1992), podocytes (Vallés-Saiz et al., 2022), pancreatic beta cells (Maj et al., 2016) as well as in peripheral neurons, including enteric neurons (Lionnet et al., 2018). Here, we show for the first time that tau is expressed in EEC, not only in the adult human colon but also in two EEC lines, namely GLUTag and NCI-H716.

Earlier studies showed that EEC exhibit neuronal features with the presence of axon-like basal processes and the expression of neuronal proteins such as synapsin 1, PGP9.5 and neurofilaments (Bohórquez et al., 2014, 2015). Our identification of tau in EEC further expands the neuronal repertoire of EEC and echoes recent publications which showed that alpha-synuclein is also expressed in EEC (Chandra et al., 2017; Casini et al., 2021; Amorim Neto et al., 2022). The observation that alpha-synuclein EEC lie in close proximity to alpha-synuclein– expressing enteric neurons led Liddle and collaborators to posit that the EEC might be critically involved in the circuit between the gut lumen and the brain for the bottom-up propagation of PD pathology (Chandra et al., 2017). Our current findings together with our previous data showing that enteric neurons express tau (Lionnet et al., 2018) suggest that such a scenario could also occur in tauopathies. It should be however borne in mind that, unlike PD and synucleinopathies (De Guilhem De Lataillade et al., 2020), all existing studies suggest that pathological tau species are not observed in the gut of subjects with either AD or PSP (Shankle et al., 1993; Lionnet et al., 2018; Dugger et al., 2019). Further studies are therefore necessary to determine if pathological tau species are present in the gut of patients with tauopathies, either in enteric nerves or EEC (Derkinderen et al., 2021). Regarding the EEC, the identification of these potential pathological tau species in the gut could greatly benefit from novel approaches, for example by combining laser capture microdissection of EEC (Blatt and Srinivasan, 2008) with ultrasensitive amplification techniques of aggregated proteins such as real-time quaking-induced conversion (Wu et al., 2022).

Six isoforms of tau are expressed in adult human brain by alternative splicing from a single gene. Regulated inclusion of exons 2 and 3 yields tau isoforms with 0, 1 or 2 N-terminal inserts (0N, 1N, 2N respectively), whereas exclusion or inclusion of exon 10 leads to expression of tau isoforms with three (3R) or four (4R) microtubule-binding repeats. The various splice combinations of tau are thus abbreviated-0N3R, 0N4R, 1N3R, 1N4R, 2N3R, 2N4R-encoding six proteins isoforms ranging from 352 to 441 amino acids in length (Goedert & Jakes, 1990). In the current study, we identified 0N3R/0N4R and 0N3R/1N3R as the two main tau isoforms expressed in GLUTag and NCI-H716 cell lines. The reasons why EEC lines only express a subset of tau isoforms remains to be determined but the observation showing that the intracellular sorting of tau in different cell compartments is isoform-dependent may provide a clue (Liu and Götz, 2013). It has been indeed reported that 0 N, 1N and 2N isoforms are primarily localized to the cell bodies, nucleus and axons, respectively (Liu and Götz, 2013), thereby suggesting that tau in EEC lines is primarily cytoplasmic. In order to properly compare the isoform profile of tau between human adult brain and EEC from human adult colon, it would have been useful to study in more detail tau isoforms in human EEC. We were however unable to do so for two main reasons: first, apart from 3R and 4R antibodies, other specific antibodies for 0N, 1N and 2N isoforms did not work well for immunohistochemistry on paraffin sections, at least in our hands; second, in contrast to our previous research on the ENS, the use of frozen colonic sections for western blot analysis was not possible because of the sparse distribution of EEC along the intestinal epithelium (Lionnet et al., 2018). Again, this precise characterization could benefit from laser microdissection approaches (Blatt and Srinivasan, 2008).

The phosphorylation of tau at multiple serine and threonine sites is the predominant mechanism by which tau functions are regulated (Guo et al., 2017). Dephosphorylation of tau from EEC lines produced a downwards shift demonstrating that EEC-tau is, like CNS tau, phosphorylated (Hanger et al., 2002). Using a phosphor-specific antibody, we further showed that tau was phosphorylated at Thr205 in GLUTag cells under basal condition and that phosphorylation at this residue was regulated by SCFA. What can be the role of tau and the consequences of tau phosphorylation from a functional point of view? In this regard, it is tempting to compare our findings to those obtained in pancreatic β-cells. EEC and pancreatic β-cells share similar pathways of differentiation during embryonic development (Ryu et al., 2018). Remarkably, like EEC, pancreatic β-cells, also express tau (Maj et al., 2016; Wijesekara et al., 2018) and several studies showed that tau is critically involved in pancreatic β-cells function, insulin secretion and glucose homeostasis (Maj et al., 2016; Wijesekara et al., 2018). One might therefore suggest that, EEC-tau similarly regulates the secretion of peptide-hormones and that such a regulation is mediated via tau phosphorylation. Further experiments performed after silencing tau in EEC will be needed to answer this question.

In conclusion, we have characterized tau in the human colon and in EEC lines and we show that EEC tau phosphorylation can be regulated by SCFA. The data we have acquired on tau in EEC strongly supports additional future studies aimed at expanding our knowledge of peripheral pathology in tauopathies and at deciphering the physiological role of tau in CEE.

## Author’s contribution

GC, NB, BC, IG, TO performed the experiments. MRD managed tissue sampling and biobanking. MRD, GLD and PD supervised the study. MN and HB provided critical feedback and helped shape the research. PD wrote the first draft of the manuscript. MRD, PD and GLD wrote the final version of the manuscript.

## Acknowledgements

We acknowledge the IBISA MicroPICell facility (Biogenouest), member of the national infrastructure France-Bioimaging supported by the French national research agency (ANR-10-INBS-04) for confocal microscopy pictures.

## Data availabilty statement

The datasets generated during and/or analysed during the current study are available from the corresponding authors on reasonable request.

